# Neuroinflammatory crosstalk between microglia and astrocytes increases viral replication in an iPSC-derived model of CNS HIV infection

**DOI:** 10.1101/2025.08.29.673049

**Authors:** James D Gesualdi, Jude Prah, Shiden Solomon, Jayden Cyrus, Ernesto Baçi, Peter J Gaskill, Çagla Akay-Espinoza, Kelly Jordan-Sciutto

## Abstract

People living with HIV suffer multiple comorbid conditions related to chronic inflammation at increased rates compared to the general population, even when on effective antiretroviral therapy. In particular, current data indicate that the increased incidence and severity of neurocognitive impairment (NCI) are associated with unresolved neuroinflammation. Attempts to treat NCI in people living with HIV by reducing inflammation have thus far been unsuccessful, suggesting that a more mechanistic understanding of inflammatory processes in the CNS during HIV is necessary. Here, we use iPSC-derived microglia (iMg) and astrocytes (iAst) to model HIV infection in the CNS. We show that our iMg robustly express markers associated with microglial identity and are susceptible to HIV infection, but exhibit lower HIV replication rates and weaker immune response to HIV challenge compared to monocyte-derived macrophages. Coculture of iAst with iMg leads to a much stronger pro-inflammatory immune response, and, surprisingly, a robust increase in rates of HIV replication. Increased replication in iMg/iAst cocultures is associated with higher levels of multiple pro-inflammatory cytokines, including TNFα, which is produced by iAst upon exposure to HIV-infected iMg. Addition of exogenous TNFα to iMg during HIV infection is also sufficient to increase rates of replication, and neutralization of TNFα via adalimumab/Humira treatment in iMg/iAst cocultures reduces replication. Blocking NF-kB signaling with iKK inhibitor Bay-11-7082 (Bay-11) demonstrates that increased HIV replication in iMg/iAst cocultures is due to increased NF-kB activity. Finally, we show that in HIV-infected iMg there is movement of lysosomes to the periphery of the cell membrane and release of lysosomal content into the extracellular space, suggesting that this dysregulated lysosomal flux could further contribute to the pro-inflammatory microenvironment. We propose that this altered lysosomal trafficking and increased cytokine production drives a pro-inflammatory phenotype in glia and represents a potential source of unresolved neuroinflammation in people living with HIV.

## Introduction

Approximately 40 million people worldwide are currently living with HIV. Since the advent of anti-retroviral therapy (ART), progression to acquired immunodeficiency syndrome (AIDS) has greatly declined^1^. Most people living with HIV (PLWH) on ART successfully achieve viral suppression if their treatment is not interrupted, eventually reaching a state in which viral RNA is undetectable in peripheral blood and transmission is not possible^2^. Indeed, PLWH on ART now have a lifespan nearly comparable to people without HIV^1,3^. Despite this remarkable progress, PLWH remain at increased risk for multiple chronic noncommunicable comorbidities, with heart disease, cancer, and cognitive decline being more likely to impact PLWH at earlier ages^4, 5^. Symptoms of neurocognitive impairment (NCI) are especially prevalent in PLWH once they reach the age of 50^4^. NCI represents a significant unmet challenge because roughly 50% of PLWH on ART develop some level of NCI or are diagnosed with HIV-associated neurocognitive disorder (HAND)^6, 7^.

HAND is a spectrum of cognitive, behavioral, and motor deficits of varying severity that are often observed in PLWH. Importantly, symptoms of NCI or HAND tend to persist despite suppressive ART^8^. Persistent NCI in the context of suppressed viremia reduces quality of life for PLWH and therefore necessitates a deeper mechanistic understanding of the underlying pathological processes in the central nervous system (CNS).

Chronic inflammation is a pathological hallmark common to the diseases and comorbidities associated with HIV^4, 9^. Increases in systemic TNFα, IL-6, and IL-1β in PLWH are associated with increased incidence of both end-organ damage^10^ and HAND^8^. Further, various single nucleotide polymorphisms in coding exons of *TNF* have been shown to increase the risk of several NCI symptoms in PLWH^11^. Additionally, indicators of neuroinflammation in the cerebrospinal fluid (CSF) such as increased levels of neopterin, CCL2, TNFα, and IL-6 are associated with an increased severity of NCI in PLWH^12, 13^. Given this association of multiple pro-inflammatory species with increased incidence and severity of comorbid conditions in PLWH, numerous immunomodulatory therapies have been explored in attempts to resolve chronic inflammation and improve quality of life for these individuals with limited success^5,14^. Therefore, a mechanistic model of sources of neuroinflammation in HAND is essential to inform clinicians of ideal cellular targets for anti-inflammatory adjunctive therapies that may be capable of mitigating NCI.

The precise mechanism through which HIV first accesses the CNS is a subject of ongoing study^15, 16, 17^. However, it is clear that myeloid cells in the CNS, including microglia and border-associated macrophages, begin harboring HIV shortly after initial infection^18^ and typically before seroconversion^19^. This suggests that myeloid cells in the CNS serves as a viral reservoir for HIV in virtually all cases. In the CNS, resident macrophages and microglia express CD4, CCR5, and CXCR4, and are the primary targets for HIV infection^20^. Even in the presence of suppressive ART, low level viral transcription and downstream neuroinflammation can occur in PLWH, and increased persistence of HIV transcription is associated with increased severity of NCI^19, 21, 22, 23^. Interestingly, the extent of neuroinflammation and neurotoxicity in PLWH on ART often exceeds the degree of pathology that can be explained by the modest levels of viral infection and HIV gene expression observed in the CNS. Indeed, recent transcriptomic studies of ex-vivo microglia isolated from PLWH on ART have shown that only roughly 0.5% of microglia harbor viral RNA transcripts^24^. This indicates that ongoing neuroinflammation and injury occur in the absence of direct viral-induced damage. In other words, there must exist mechanisms that contribute to tissue damage in the CNS that are at least somewhat independent of ongoing productive viral replication. These findings have prompted multiple studies on the role that bystander cells such as astrocytes may play in the context of HAND and HIV NCI.

Astrocytes, the most numerous cell type in the CNS, perform many critical neurotrophic functions under homeostatic conditions^25^. Additionally, astrocytes act to maintain the integrity of the blood brain barrier and contribute to immune responses to potential infectious challenges in the CNS^25^. Some studies suggest that astrocytes are capable of harboring HIV infection^26, 27^, but evidence of active viral replication in astrocytes is limited and this remains an area of active debate^19, 26, 27, 28^. However, evidence of reactive or gliotic phenotypes among astrocytes is a common pathologic hallmark of many neuroinflammatory and neurodegenerative conditions and is associated with more severe NCI in HAND^19, 29, 30^, highlighting the role for astrocytes in HIV neuropathogenesis.

Studying the immune response of both microglia and astrocytes during HIV infection of the CNS at the mechanistic level has been challenging due to the difficult nature of obtaining and culturing microglia *in vitro* without incurring broad loss of their *in vivo* characteristics^31^. Here, we use a coculture model of induced-pluripotent stem cell (iPSC)-derived microglia like cells (iMg) and astrocytes (iAst) to study the immune dynamics and crosstalk between these two glial populations in the context of HIV infection. We show that iMg express critical markers of microglial lineage and function and are susceptible to HIV infection. We also demonstrate that HIV is capable of evading various canonical anti-viral immune responses in iMg and, surprisingly, that the coculture of iAst with HIV-infected iMg robustly increases the rate of viral replication. Our analyses reveal that this phenotype in cocultures is associated with the presence of pro-inflammatory cytokines produced by iAst, which in turn drive NF-kB activation in iMg, accelerating viral replication. Targeting pro-inflammatory cytokine production via neutralizing antibody blockade or attenuating NF-kB signaling via small molecule inhibition significantly reduces HIV replication rates observed in the iMg/iAst coculture model. Finally, we demonstrate that HIV infection in iMg leads to repositioning of lysosomes to the cell periphery and to increased exocytosis of lysosomal content into the extracellular space. This process suggests a potential microglia-dependent mechanism through which iAst become gliotic and begin producing TNFα and other pro-inflammatory molecules, driving the increased HIV replication observed in cocultures. Our findings provide a mechanistic foundation for the neuroinflammation driven injury in PLWH and suggest potential therapeutic avenues for reducing unresolved pathologic inflammation despite effective ART.

## Materials and Methods

### Reagents

**Table 1.**
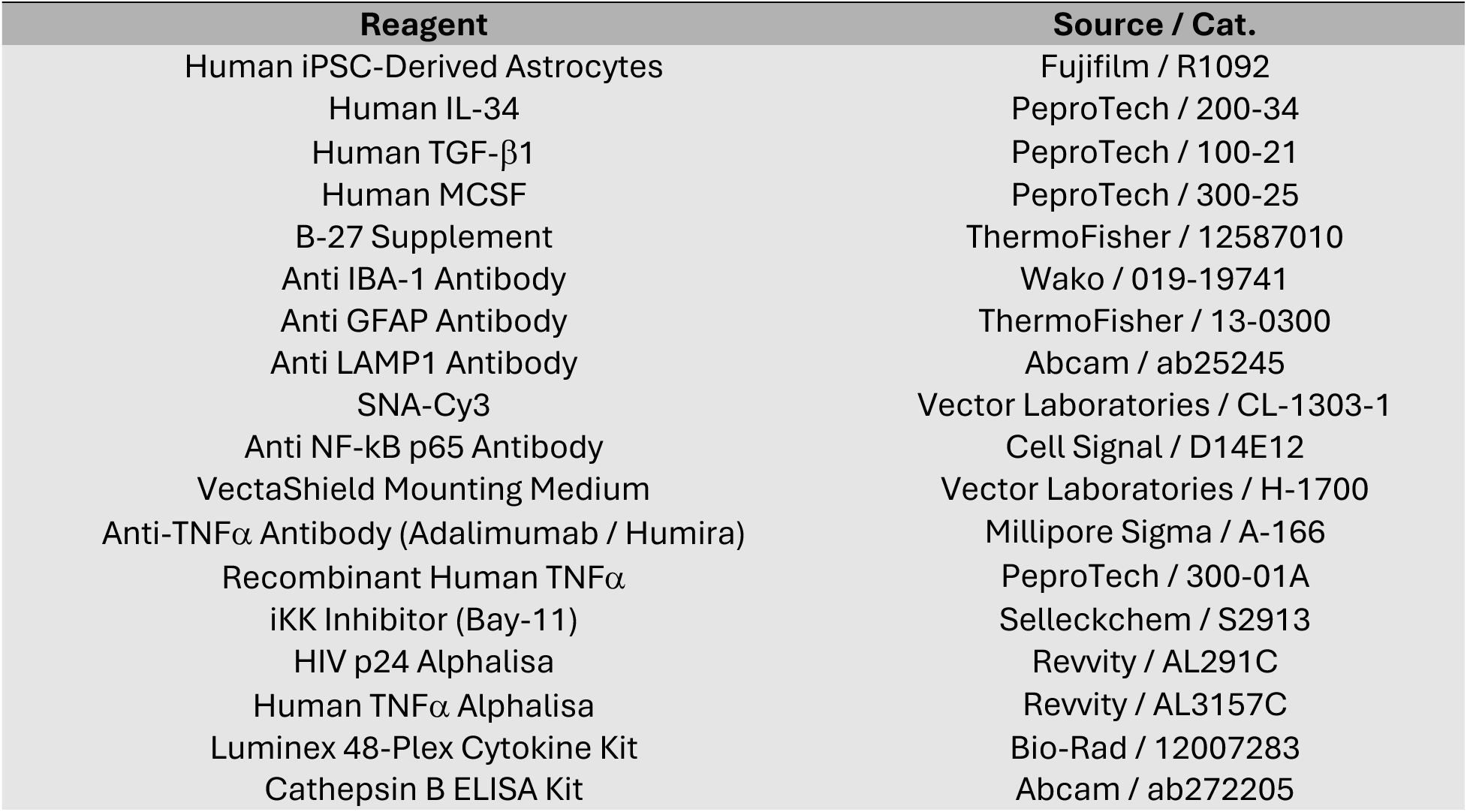
Reagents used in these studies.

### Differentiation iMg and monocyte-derived macrophages

Human pluripotent stem cells derived from the skin fibroblasts of healthy adult donors were differentiated into common myeloid progenitors (CMPs), as previously described^32, 33^. CMPs purchased from the Human Pluripotent Stem Cell Core at the Children’s Hospital of Philadelphia were cultured in RPMI (HyClone) containing 10% FBS containing penicillin/streptomycin with 100 ng/mL IL-34, 50 ng/mL transforming growth factor beta TGFβ, and 25 ng/mL macrophage colony stimulating factor (MCSF) for 11 days, as previously described^34^. CMPs were plated at a density of 5 ^x^ 10^5^ cells /mL for all experiments, either in 24 or 96-well glass-like polymer coated culture plates (Cellvis). For immunostaining on glass coverslips, iMg were plated and differentiated at the same density in 24 well glass-like polymer coated plates with circular coverslip inserts.

Monocyte-derived macrophages (MDMs) were differentiated from peripheral blood monocytes isolated from healthy volunteers using magnetic bead-based separation carried out by the University of Pennsylvania Human Immunology Core, as previously described^35^. Monocytes were plated at 5 ^x^ 10^5^ cells /mL and differentiated in DMEM (Thermo Fisher) containing 10% FBS with penicillin/streptomycin and 20 ng/mL granulocyte-macrophage colony-stimulating factor for seven days, as previously described^35^.

### iMg/iAst cocultures

Fully differentiated iAst were obtained from Fujifilm^36^. For monocultures of iAst and cocultures of iMg/iAst, glass-like polymer coated culture plates (CellVis) were coated with 1.2 ng/mL laminin in phosphate-buffered saline (PBS) for 2h. iAst were collected in BrainPhys (Thermo Fisher) supplemented with B27 (Thermo Fisher) and plated at a density of 10^5^ cells/mL. For iMg/iAst cocultures, CMPs were plated at 5 ^x^ 10^5^ cells / mL in wells containing 10^4^ iAst. iMg/iAst cocultures were maintained in iMg media containing 10% FBS, 1X penicillin/streptomycin with 100 ng/mL IL-34, 50 ng/mL TGFβ, and 25 ng/mL M-CSF in addition to B27.

### Adalimumab, TNFα, and Bay-11 formulation

Adalimumab (Millipore Sigma), recombinant human TNFα (PeproTech), and Bay-11 (Selleckchem) were formulated in anhydrous DMSO. Working stocks were diluted in sterile PBS before preparing media aliquots with appropriate drug concentrations for treatment of monocultures or cocultures during HIV infection.

### HIV infection

For all experiments except RNA sequencing, fully differentiated iMg, MDMs, and iMg/iAst cocultures were treated with 1 ng/mL HIV_ADA_ for 24 hours. In all experiments, a heat inactivated (dI) viral stock was included as a mock infection. After 24 hours of incubation, excess virions were removed via three sequential 60% washes with RPMI on post-infection day 0 (PID. 0). Supernatants were collected from cultures at 3, 6, 9, or 12 days post-infection (PID 3, 6, 9, or 12) for various analyses. Supernatant collection was accompanied by 50% media changes on infected cells. For experiments in which infected cultures were treated with adalimumab, recombinant TNFα, or Bay-11, the reagents were diluted to the indicated concentrations in the appropriate media for each culture condition and used to replace the media after supernatant collection beginning on PID3. For immunostaining, at indicated timepoints, the cultures were fixed using 4% paraformaldehyde (Thermo Fisher) in PBS for 20 min at room temperature.

### RNA isolation and sequencing

Fully differentiated iMg and MDM cultures were infected with 50 ng/mL HIV Jago, using a higher concentration of HIV to increase the number of infected cells as require for sequencing. Following supernatant collection on PID 12, MDM and iMg were lysed in wells using the RLT buffer (Qiagen RNeasy Mini Kit 74104), and pooled RNA was isolated according to manufacturer protocols. RNA purity and concentration were confirmed using Nanodrop, and Zymo RNA Clean & Concentrator-25 was used on any samples that did not pass initial quality control based on an absorbance of ≥ 2 at 260/280 nm and an absorbance of ≥1.8 at 260/230 nm, with minimum concentration of 50 ng/uL.

The remaining steps were performed by Azenta Life Sciences. Briefly, purified RNA was used for mRNA-enriched library preparation and sequencing via Illumina HiSeq, with an average read depth of 22 ^x^ 10^6^ reads/sample. Reads were trimmed using Trimmomatic v.0.36 and mapped to the *Homo Sapiens* GRCh38 reference genome using STAR aligner v.2.5.2b. Next, in the R computing environment, raw gene counts were calculated using featureCounts from the Subread package v.1.5.2. Raw counts were read into edgeR v.4.4, and low counts below 1 counts/million were filtered out as unexpressed genes. Counts were then normalized as transcripts/million^37^. Significantly differentially expressed genes (DEGs) were defined as those with a log-fold change ≥ 1.0 and an adjusted p value of ≤ 0.05. DEGs were analyzed using the gene set enrichment analysis package GSVA v.3.2^38^. Raw data from the bulk RNA sequencing experiment are available at NCBI GEO (accession GSE143685).

### Quantification of HIV p24 and cytokines in culture supernatants

Supernatants from monocultures of iMg, MDMs, iAst, and from cocultures of iMg/iAst were collected following HIV infection and various drug treatments at PID 3, 6, 9, or 12. Supernatants were analyzed in duplicate by AlphaLISA to determine the levels of HIV capsid protein p24 (Revvity AL291C) or TNFα (Revvity AL3157C), as previously described according to manufacturer’s protocols^39^. Briefly, supernatants were incubated with an anti-target primary antibody conjugated to streptavidin and biotinylated fluorescent acceptor beads for 1 h at RT. Next, fluorescent donor beads were added to the samples, which were incubated for 30 minutes at RT. The fluorescence was quantified using an Envision Excite plate reader and raw data were interpolated using a standard curve to quantify the concentration of target analytes. The limits of detection on these assays were 1.8 pg/mL (p24) and 1.41 pg/mL (TNFα).

For unbiased cytokine assays, supernatants were assayed in duplicate using the Bio-Plex Pro Human Cytokine Screening Panel, 48-Plex (Bio Rad #12007283) by the University of Pennsylvania Human Immunology Core. Raw pg/mL values were presented indicated in scatter plots and fold change normalized to negative control (iMg treated with dI HIV) were presented as heat maps.

### iAst Monoculture Treatments

iAst monocultures were plated at 10^4^ cells per well on 96-well glass-like polymer coated plates coated with 1.2 ng/mL laminin for 2h at 37°C. iAst were treated with either 50% conditioned media from HIV-infected iMg, 100 ng/mL lipopolysaccharide, 10 ng/mL IL-1ß, or 10 ng/mL HIV_ADA_ for 24 h. TNFα levels in collected supernatants were analyzed by alphalisa, or Bio-Plex Pro Human Cytokine Screening Panel, 48-Plex. Following supernatant collection, cells were fixed using 4% paraformaldehyde in PBS for 20 min at room temperature for immunostaining.

### Immunofluorescence staining

Cells were fixed in culture plates with 4% paraformaldehydein PBS for 20 min and washed with PBS and PBS containing 0.1% Tween-20 (PBS-t) (Thermo Fisher). Cells were then blocked and permeabilized with 1.0% BSA in PBS containing 0.1% Triton-X (Thermo Fisher) (PBS-Tx) for 30 min. Cells were then incubated overnight at 4C with primary antibodies targeting cell-surface markers (IBA-1 or SNA for iMg, GFAP for iAst) and proteins of interest (LAMP1, NF-kB p65) at appropriate dilutions in blocking buffer. Cells were then incubated for 2 h at RT with fluorescently labeled secondary antibodies corresponding to appropriate antibody host isotypes (1:200) and 4′,6-diamidino-2-phenylindole, dihydrochloride (DAPI, 1:1000). Stained cells were washed with PBS and stored in PBS at 4C until imaging. For iMg differentiated on glass coverslips, coverslips were removed from culture plates with forceps and then mounted on slides using 20uL of VectaShield antifade mounting medium.

Images were captured using a Keyence BZ-X710 widefield fluorescent microscope with a 20X objective using DAPI, FITC, and Cy3 filter cubes or Leica TCS SP8 STED Confocal with a 40X objective. Scale bars were applied using ImageJ annotation software. For LAMP1 images, lysosomes were quantified near the cell membrane by manually defining a region of interest within ten pixels of the cell membrane as indicated by IBA-1 staining and then measuring the strength of local LAMP1 staining by quantifying the average intensity of 488 staining. For NF-kB p65 quantification, regions of interest were automatically selected via thresholding whole cells and nuclei of microglia as indicated by SNA and DAPI staining, respectively, and strength of local NF-kB p65 signal was calculated by measuring the intensity of Cy5 staining in the indicated regions of interest.

### Statistical analysis

Bulk RNA sequencing included three MDM donors and three iMg lines. For supernatant analyses and immunostaining, 2-3 differentiations of up to four iMg lines were used, with data averaged across differentiations to ensure that the observed changes were biologically and technically consistent. HIV p24 and cytokine analyses from culture supernatants were evaluated using two-way ANOVA or multiple paired t-tests to compare across culture conditions and treatments, depending on the experimental design. Whenever possible, data are visualized as a ‘SuperPlot’ in which technical replicates within a biological replicate are color coded, and averages among the constitutive technical replicates for each biological replicate are overlayed as a solid-colored point^40^. ANOVA and t-tests are performed on biological replicates.

## Results

### iMicroglia express canonical cell-surface markers and are susceptible to HIV infection

In a previous study^33^, iMg differentiated using this cytokine cocktail have been shown to successfully exhibit a ramified morphology (Figure 1a) and robustly express classic microglial lineage markers, including *AIF1*, *TMEM119*, *P2RY12*, *SALL1*, *SMAD4*, *PTPRC*, *ITGAM*, *TGFBR1*, and *TGRBR2*^41^ (Figure 1b). We have also previously reported that these iMg exhibit synaptophagocytic function^34^, replicating key *in vivo* characteristics of human microglia. iMg also expressed high levels of HIV-associated cell-surface receptors *CD4*, *CCR5*, and *CXCR4* (Supplementary Figure 1a). Accordingly, HIV_ADA_ was capable of entry and active replication in iMg. As expected, replication rates in iMg are reduced compared to those observed in human MDMs, suggesting that microglia are a suboptimal cellular host for HIV replication (Figure 1c), despite being permissive to infection. This may be due to a reduced expression of the HIV coreceptor *CCR5* in iMg compared to the MDMs (Supplementary Figure 1a). These combined traits of infectability with reduced virion production are consistent with features of a cellular reservoir.

**Figure 1:**
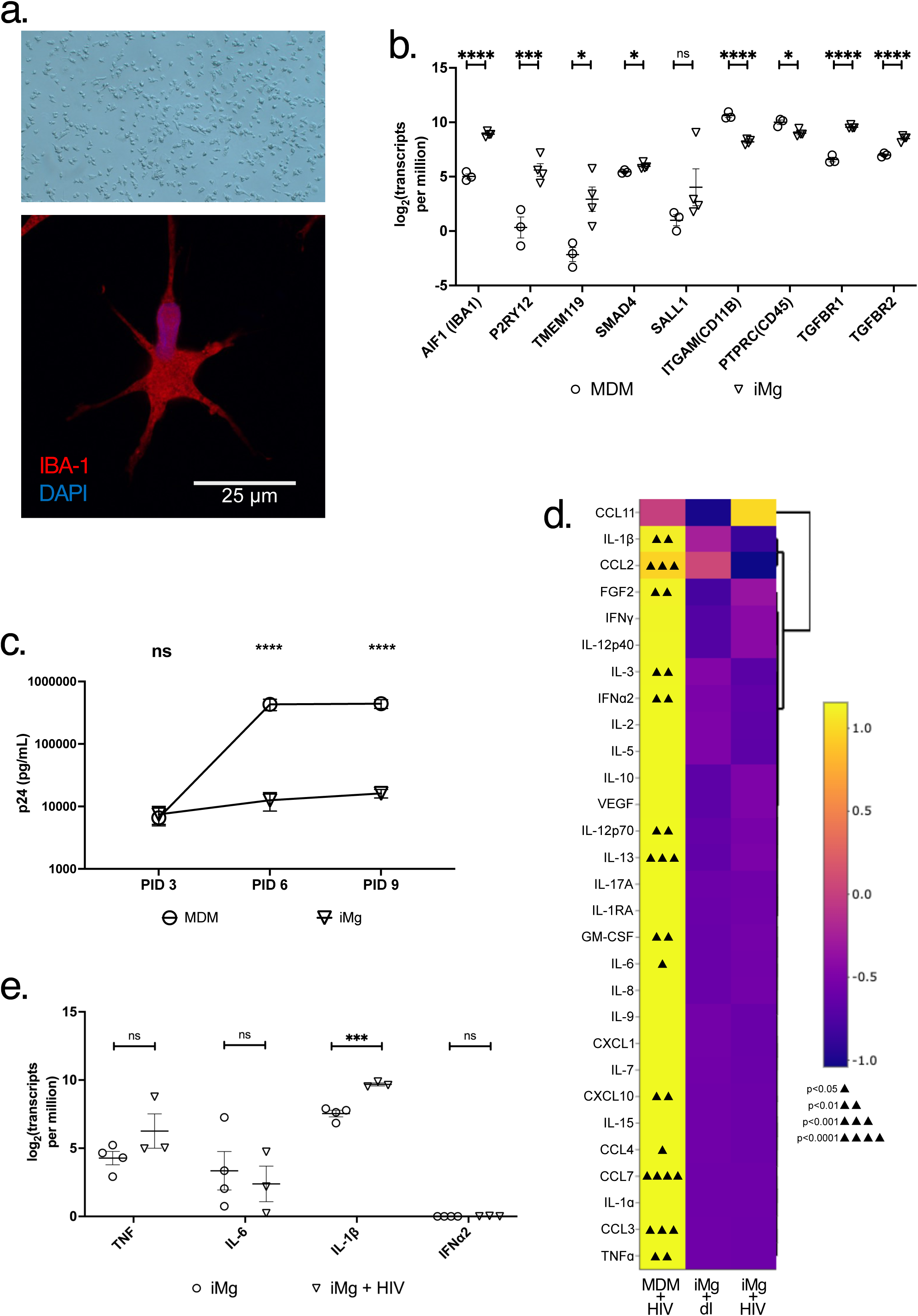
iMicroglia express canonical cell-surface markers and are susceptible to HIV infection. (a). iMicroglia at 11 days *in vitro* via brightfield (top, 10X) or confocal (63X, bottom) microscopy. (b). Expression of indicated microglial lineage genes in iMg at 11 DIV compared to expression in MDM in log transformed transcripts per million. (c). HIV replication in iMg compared to MDM as measured by levels of HIV capsid protein p24 via alphalisa following infection with 1 ng/mL HIV_ADA_. Data are expressed as Mean and SEM of 2 – 3 biological replicates in duplicate. (d). Heatmap indicating relative levels of indicated cytokines and chemokines produced by HIV-infected MDM (left) or iMg (right) compared to mock-infected iMg (center) at the protein level. Significant upregulation of analytes compared to mock-infected iMg are annotated via triangles in individual cells. Data are condensed from 2(MDM) to 5 (iMg) biological replicates. (e). Expression of indicated cytokines in iMg with or without HIV infection at PID 12 in log transformed transcripts per million * = p < 0.05, *** = p < 0.001, **** = p < 0.0001. Multiple paired T-tests.

MDMs have been shown to produce a robust pro-inflammatory response, including inflammasome activation, following HIV infection^42, 43, 35^. Accordingly, we found that HIV-infected MDMs secreted pro-inflammatory cytokines, such as IL-1β, TNFα, and IL-6 and viral restriction factors such as IFNα2 (Figure 1d). In contrast, iMg failed to secrete any of these pro-inflammatory molecules at significant rates following infection with HIV at either the protein or the mRNA level (Figure 1d, 1e), with the exception of IL-1β, which was observed only at the transcript level(Figure 1e) indicating that mature IL-1β cleavage and secretion was not occurring in HIV-infected iMg monocultures, indicating reduced canonical anti-viral and pro-inflammatory immune responses in iMg compared to MDMs.

### Coculture of iMg with iAst leads to increased rates of HIV replication

Given the weak immune response of iMg following challenge with HIV, we hypothesized that including iAst in coculture would enable a more pronounced immune activation. Various groups have also shown that differentiating iMg in the presence of other neuronal cells produces a more functional microglia population with a morphology more akin to homeostatic ramification typically observed *in vivo^44^*. Similarly, our iMg adopted a more readily observable ramified morphology under homeostatic conditions when differentiated in the presence of iAst (Figure 2a).

**Figure 2:**
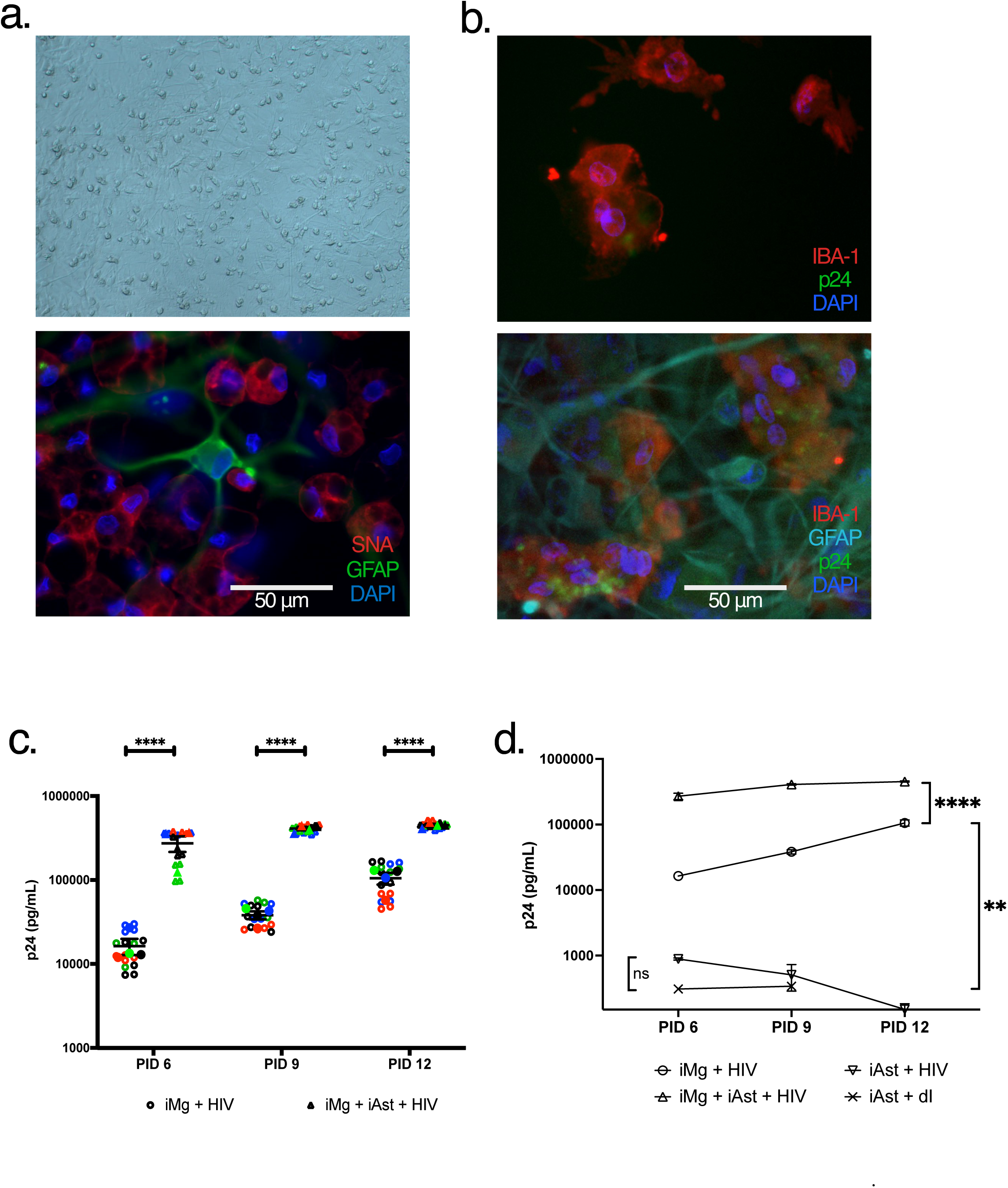
Coculture of iMicroglia with iAstrocytes leads to increased rates of HIV infection. (a). iMg cocultured with iAst at 11 days in vitro via brightfield (top, 10X) or widefield (bottom, 40X) microscopy. (b). Widefield image of iMg (top) or iMg/iAst (bottom) cocultures following HIV infection captured at 40X at PID12. (c). HIV replication rates measured by HIV capsid protein p24 in iMg monocultures (circles) or iMg/iAst coclutures (triangles) after infection with 1 ng/mL HIV_ADA_. In panels b and c, distinct biological replicates are indicated by different colors, and averages + SEM of technical replicates within each of the groups of biological replicates are indicated by the overlayed solid-colored points and error bars. (d). HIV replication rates measured by HIV capsid protein p24 in iMg monocultures (circles), iMg + iAst cocultures (triangles), iAst monocultures (inverted triangles) after infection with 1 ng/mL HIV_ADA_ or 1 ng/mL heat inactivated (dI) HIV (x). iMg groups include data condensed from four biological replicates, iAst groups include data condensed from three technical replicates. * = p < 0.05, ** = p < 0.01, **** = p < 0.0001. Two-way ANOVA (c) or Multiple paired T-tests (d).

Surprisingly, the coculture of iMg with iAst led to a significant increase in HIV replication. Presence of iAst in a 1:5 ratio with iMg produced HIV replication an order of magnitude higher than iMg alone, as measured by p24 Gag(p24) levels in culture supernatants (Figure 2c). Importantly, this difference was not observed due to HIV replication in iAst themselves, as p24 was not observed via immunostaining in iAst in cocultures (Fig 2b) and replication did not occur to any appreciable level in iAst monocultures (Figure 2d). This finding suggested that iAst increased the rate of active HIV replication in iMg without acting as a direct cellular target for HIV in our model. We next profiled the immune response in iMg/iAst cocultures to identify potential differences associated with this increase in viral replication.

### iAst adopt a pro-inflammatory disease-associated phenotype when exposed to HIV-infected iMicroglia

Based on the impact of iAst on HIV replication rates in iMg, we evaluated the cytokine profile of HIV-infected iMg in the presence and absence of iAst. HIV-infected iMg in monocultures exhibited a modest immune response marked by the production of CCL2 and IL-8 (Figure 3a, 3b). In contrast, HIV-infected iMg cocultured with iAst showed significantly increased production of CCL2, VEGF, IL-6, and TNFα suggesting a more robust pro-inflammatory immune response to viral challenge (Figure 3a-3c). Interestingly, IFNα2 was not significantly upregulated in iMg/iAst cocultures compared to iMg monocultures, suggesting that the nature of the immune response produced by cocultures in response to HIV infection was more broadly pro-inflammatory rather than specifically anti-viral in nature.

**Figure 3:**
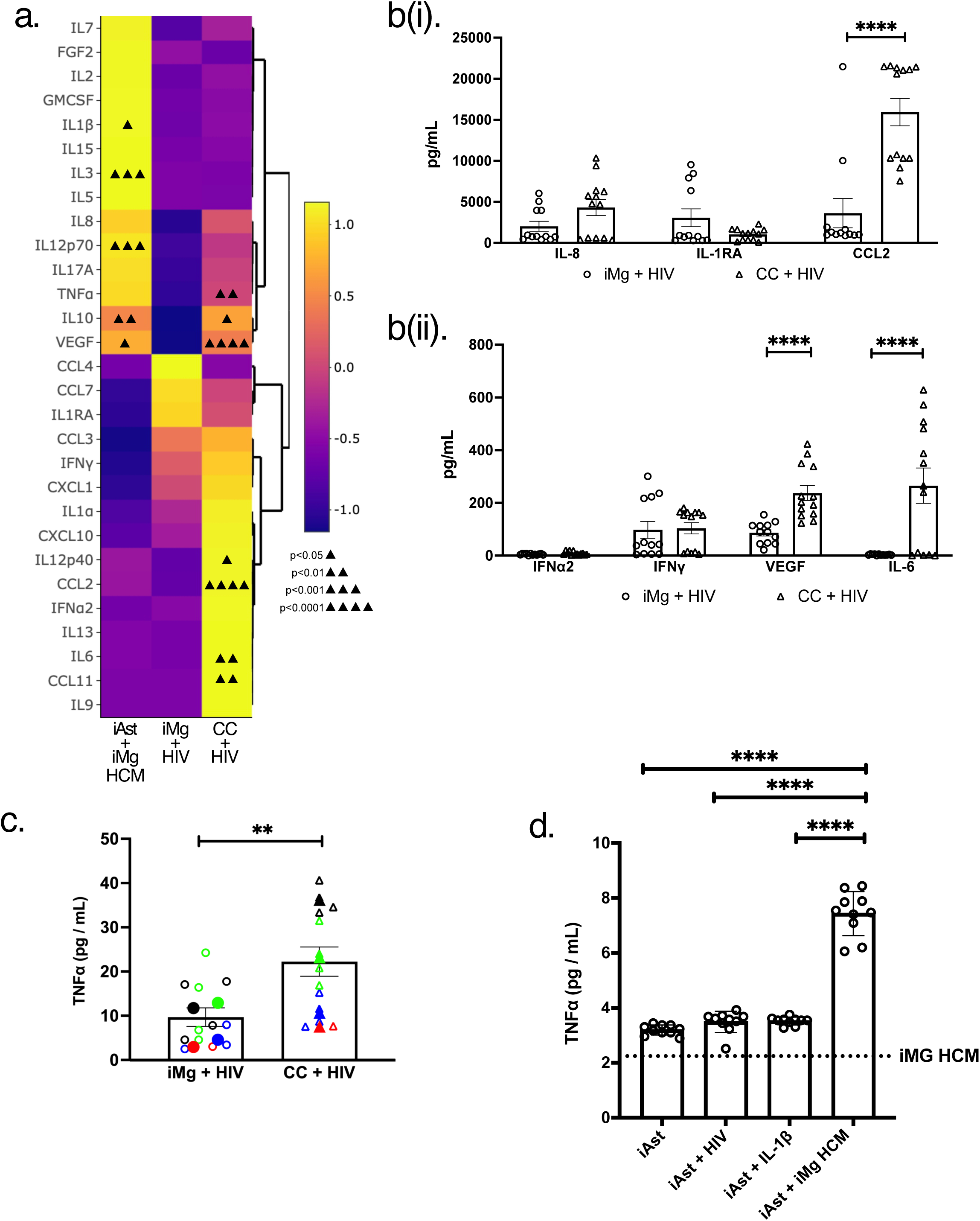
iAstrocytes adopt a pro-inflammatory disease-associated phenotype when exposed to HIV-infected iMicroglia. (a). Heatmap indicating relative levels of indicated cytokines and chemokines secreted by HIV-infected iMg monocultures (iMg+HIV, center), iMg/iAst cocultures (CC+HIV, right), and monocultures of iAst exposed to supernatants from HIV-infected iMg (iAst+iMg HCM, left), measured by biorad Luminex assay. Significant up/downregulation of analytes compared to HIV-infected iMg monocultures are annotated via triangles in individual cells. Data are averaged from 3 (iAst) to 5 (iMg and coculture) biological replicates (b). Raw pg/mL values measured by biorad Luminex of indicated cytokines and chemokines produced by HIV-infected iMg monocultures (circles), HIV-infected iMg + iAst coclutures (triangles). Data include 5 biological replicates. (c). TNFα levels secreted by HIV-infected iMg and HIV-infected iMg + iAst cocultures measured by biorad Luminex assay. Technical replicates within a group of biological replicates are indicated by distinct colors, and averages from a given biological replicate are expressed as the overlaid solid-colored symbols. Data include 2-4 biological replicates. (d). TNFα production measured via alphalisa for monocultures of iAst treated for 24h with the indicated stimuli. Data include 5 biological replicates per group measured in duplicate. * = p < 0.05, *** = p < 0.001, **** = p < 0.0001. Multiple paired T-tests.

Next, we measured cytokine production by iAst in monocultures exposed to conditioned media of the HIV-infected iMg to determine the pro-inflammatory molecules specifically produced by iAst (Figure 3a, 3d). The treatment of iAst with conditioned media from the HIV-infected iMg led to the production of TNFα, but not that of IL-6 (Figure 3d, Supplementary Figure 3a). This indicates that factors secreted from HIV-infected iMg were sufficient to drive astrogliosis and pro-inflammatory cytokine production in iAst. Indeed, exposure of iAst monocultures to the supernatants from HIV-infected iMg led to a pro-inflammatory phenotype in iAst that is similar to the disease-associated astrocyte state observed in several neuropathic conditions^45^.

### Astrocyte derived TNFα increases HIV replication rates in iMicroglia via NF-kB activation

TNFα has been shown to be sufficient to increase transcription of integrated HIV genomes in reservoir studies and is considered a stimulus capable of accelerating active rather than latent or transcriptionally silent HIV infection^46, 47, 48^. Indeed, TNFα was shown to be sufficient to trigger reactivation of transcriptionally silent HIV infection in a microglia-derived cell line^49^. As such, we hypothesized that iAst-derived TNFα led to an increase in HIV replication observed in iMg cocultured with iAst. We tested this via addition of exogenous recombinant TNFα to monocultures of iMg infected with HIV and antibody blockade of TNFα in infected iMg/iAst cocultures. As expected, recombinant TNFα was sufficient to significantly increase HIV replication rates in monocultures of iMg (Figure 4a). Conversely, the neutralization of TNFα in infected cocultures via adalimumab treatment significantly reduced HIV replication, but not to the level observed in untreated iMg monocultures (Figure 4b). This suggested that while TNFα could drive increased HIV replication, it is likely not the only pro-viral factor secreted by iAst that acts to promote HIV replication in iMg in our model.

**Figure 4:**
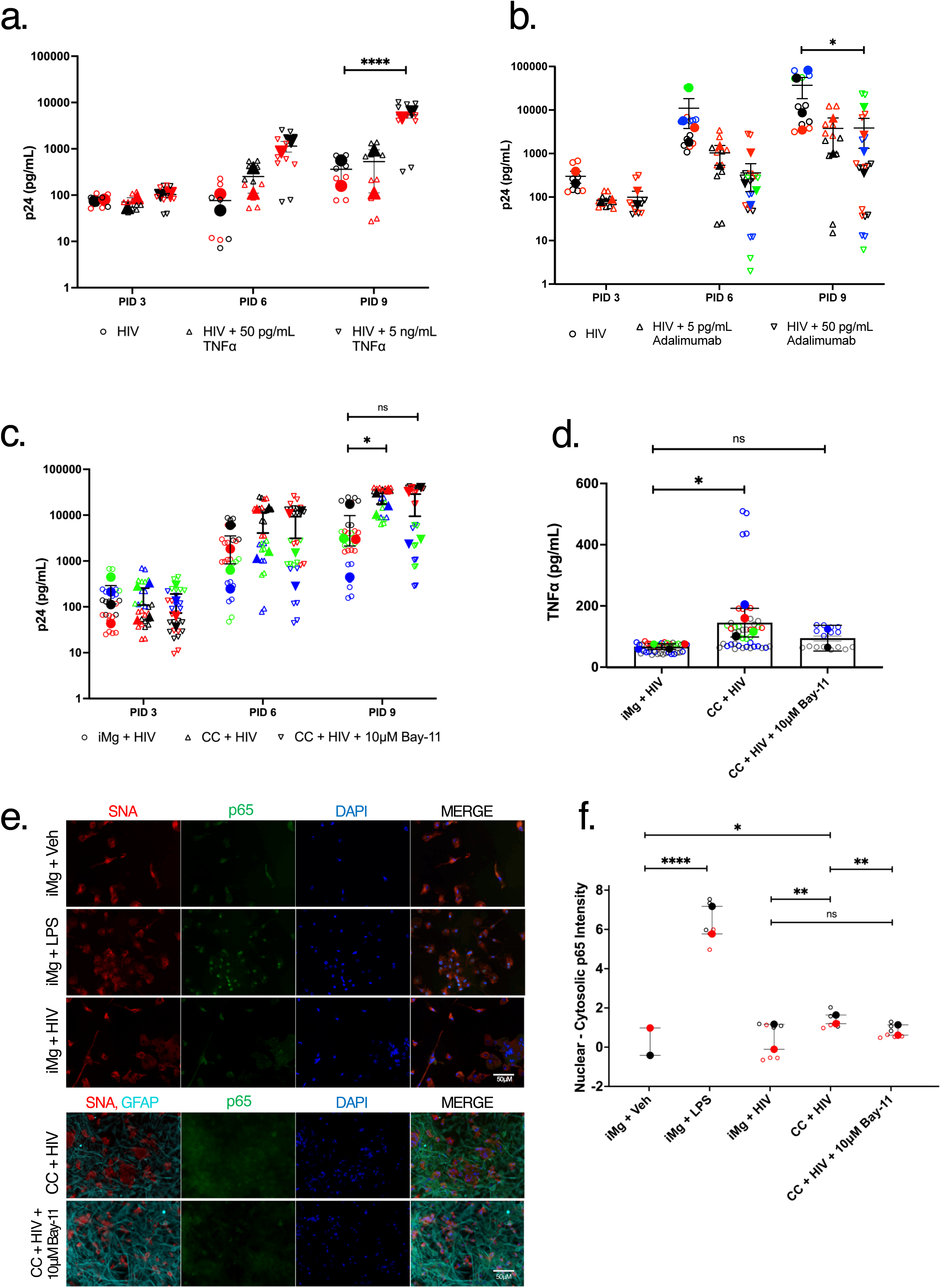
Astrocyte derived TNFα increase HIV replication rates in iMg via NF-kB activation. (a). HIV replication rates measured by HIV capsid protein p24 in iMg monocultures following infection with 1 ng/mL HIV_ADA_ and addition of recombinant TNFɑ at indicated concentrations. (b). HIV replication rates measured by HIV capsid protein p24 in iMg/iAst cocultures following infection with 1 ng/mL HIV_ADA_ and addition of TNFɑ neutralizing antibody adalimumab at indicated concentrations. (c). HIV replication rates measured by HIV capsid protein p24 in iMg/iAst cocultures following infection with 1 ng/mL HIV_ADA_ and addition of iKKɑ inhibitor Bay-11 at 10 µM. (d). TNFɑ production as measured by alphalisa in HIV-infected iMg monocultures, iMg/iAst cocultures, and iMg/iAst cocultures treated with 10 µM Bay-11. In a-d, data are representative of 2(a,d) to 4(b, c) biological replicates. Technical replicates within a group of biological replicates are indicated by distinct colors. (e). Widefield microscopy images of iMg or iMg/iAst cocultures following the indicated treatments. SNA is a lectin staining marking iMg cellular membranes in Red, GFAP shows iAst in Cyan, NF-kB p65 is indicated in green and DAPI in blue. All images captured at 40X. (f). Quantification of (e) via automatic quantification of p65 staining in nuclei subtracted from whole cell p65 staining. * = p < 0.05, ** = p < 0.01, **** = p < 0.0001. Two-Way ANOVA (a – d), Multiple paired T-tests (f).

We hypothesized that the mechanism through which TNFα augmented HIV replication was dependent of NF-kB nuclear translocation. TNFα is a known inducer of NF-kB signaling^50, 48^, and numerous studies have shown that NF-kB nuclear translocation can lead to increased transcription of the integrated HIV genome^51, 49^, thereby increasing viral replication. We tested this by treating HIV-infected iMg/iAst cocultures with the Bay-11-7082(Bay-11), which binds and inhibits inhibitory kappa beta kinase (iKK), preventing the phosphorylation and subsequent nuclear translocation of NF-kB p65^52^. Bay-11 treatment at 10 μM abrogated the observed increase in viral replication in iMg/iAst cocultures, suggesting that NF-kB signaling is critical for HIV replication in iMg (Figure 4c). Importantly, treatment of infected iMg/iAst cocultures with 10 µM Bay-11 reduced HIV replication to levels comparable to those observed in untreated iMg monocultures, suggesting that NF-kB signaling is the main mechanism driving the observed increase in viral replication in iMg/iAst cocultures. Additionally, Bay-11 treatment also abrogated the observed increase in TNFα production detected in HIV-infected cocultures, suggesting that astrocytic production of TNFα in iMg/iAst cocultures also depends on NF-kB signaling. To confirm that Bay-11 treatment successfully inhibited NF-kB dependent transcriptional regulation, we stained HIV-infected iMg monocultures and iMg/iAst cocultures with for NF-kB p65 (Fig 4e-f). Robust nuclear translocation was observed in iMg treated with 100ng/mL LPS as a positive control, showing the efficacy of this experimental design. As expected, p65 signal remained largely cytosolic in uninfected iMg, and nuclear p65 staining was more intense in HIV-infected iMg/iAst cocultures compared to HIV-infected iMg monocultures. This increase in nuclear translocation of NF-kB p65 was inhibited to the level seen in HIV-infected iMg monocultures in the presence of Bay-11.

### HIV infection causes lysosomal exocytosis in iMicroglia

Given the robust pro-inflammatory phenotype observed in iAst exposed to conditioned media from infected iMg (Figure 3b), we hypothesized that factors secreted by infected iMg following HIV infection caused reactive gliosis in iAst. A recent proteomic study of HIV-infected macrophages showed that infection led to exocytosis of lysosomes, creating a pro-inflammatory microenvironment^53^. Additionally, multiple HIV viral proteins have been shown to be sufficient to alter lysosomal position, pH, and function in various cell types^54, 55^.

Thus, we hypothesized that HIV infection in iMg would alter lysosomal distribution, leading to lysosomal exocytosis and reactivity in iAst. Indeed, monocultures of HIV-infected iMg have lysosomes more proximal to the plasma membrane as determined by LAMP1 immunostaining (Figure 5a, 5b). Additionally, supernatants from HIV-infected iMg monocultures and iMg/iAst cocultures both had increased extracellular levels of cathepsin B compared to uninfected controls, indicating the release of lysosomal content into the extracellular space following HIV infection (Figure 5c). Interestingly, both iMg monocultures and iMg/iAst cocultures showed substantial cathepsin B release upon treatment with heat-inactivated (dI) HIV_ADA_, suggesting that residual viral protein or nucleic acids remaining in this formulation are sufficient to perturb lysosomal flux in glia (Fig S5a). These data provide a mechanism through which HIV infection in iMg creates a pro-inflammatory micro-environment driving astrocytic reactivity.

**Figure 5:**
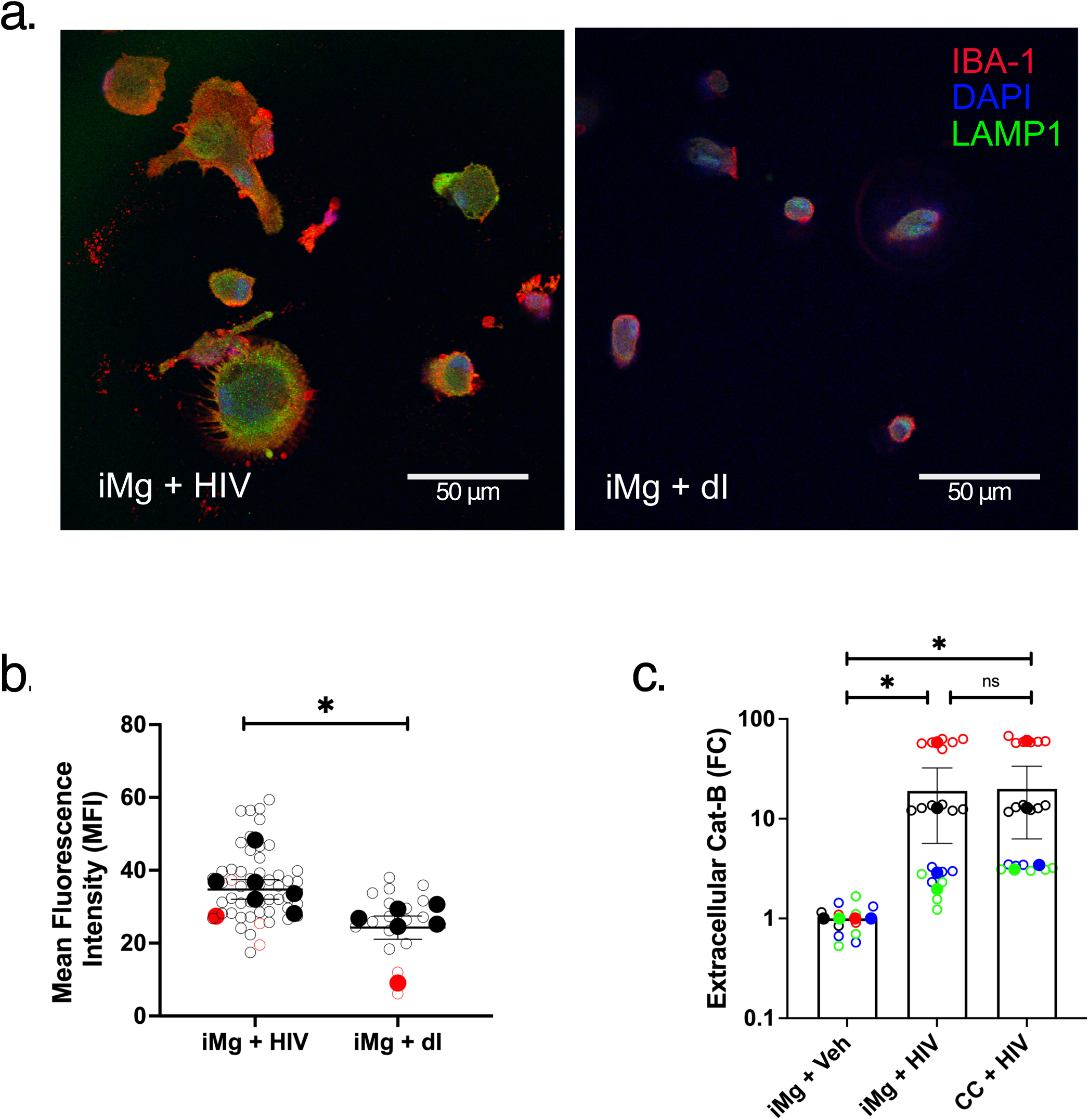
HIV infection leads to lysosomal release by iMg. (a). IBA1 (red), LAMP1 (green), and DAPI (blue) staining in iMg with (left) or without HIV infection (right) captured at 40X via confocal microscopy. (b). Quantification of (a). A 10 pixel-wide region of interest was manually drawn at the plasma membrane of each cell, and mean fluorescence intensity of LAMP1 staining was measured, indicating trafficking to the cell periphery. Distinct biological replicates are indicated by different colors, and averages + SEM of technical replicates within each of the groups of biological replicates are indicated by the overlayed solid-colored points and error bars. Data are condensed from 3 (dI) to 6 (HIV) independent experiments. (c). Cathepsin B measured in culture supernatants from indicated groups at PID 12. Data are condensed from four biological replicates and plotted as fold-change vs the negative control group, iMg + Veh. * = p < 0.05, **** = p < 0.0001. Multiple paired T-tests (b), Mann-Whitney test (c).

## Discussion

We show that our iMg reliably model *in vivo* microglia, including canonical cell surface marker expression and morphology. iMg allow the modeling of HIV infection *in vitro* and show a reduced immune response to HIV compared to other cell types susceptible to HIV infection, as shown in other recent studies of iPSC-derived microglia^56^. We also demonstrate that coculture of iMg with iAst leads to robust increases in the rate of HIV replication. This increase in viral replication is associated with astrocytic production of pro-inflammatory cytokines, particularly TNFα, which drive NF-kB signaling, in turn leading to greater rates of viral replication in iMg. We also show that iAst exposed to infected iMg or soluble factors produced by infected iMg develop a pro-inflammatory disease-associated phenotype. This astrocytic activation is associated with dysregulated lysosomal flux and increased lysosomal exocytosis from iMg following HIV infection as measured by cathepsin B release.

Recent studies on HIV infection in the CNS of PLWH on ART have shown that, while pathologic changes, such as inflammation, synaptic damage, and white matter abnormalities may be somewhat widespread^57^, productively infected leukocytes are quite rare in the CNS^58, 30, 26^. In fact, a recent study estimated that only roughly 0.5% of microglia harbor viral RNAs, indicating productive infection^24^. Despite the minimal presence of HIV virions in the CNS, neuroinflammation and CNS injury continue to accumulate overtime, leading to the onset of or worsening of NCI^12, 59, 60, 30^. The persistence of pathology in the context of a relative lack of viral infection suggests triggers of damaging neuroinflammation and gliosis that persist independently of replication^24, 61^.

Our data show that even microglia that are not actively transcribing HIV may still exhibit dysregulated lysosomal flux (Figure S5a). This may create a pro-inflammatory microenvironment due to the exocytosis of undegraded cellular materials, leading to astrogliosis. This is a potential mechanism through which neuronal injury may continue to persist and perhaps worsen in the setting of suppressive ART. The incidence of dysregulated lysosomal flux in the CNS of PLWH has not been demonstrated via immunohistochemistry. However, studies on peripheral biomarkers in Alzheimer’s Disease patients have shown increased levels of proteins associated with aberrant autophagy and lysosomal function such as cathepsin D in peripheral blood^62^. Given the abundance of studies demonstrating the ability of HIV accessory genes to alter lysosomal distribution and drive lysosomal exocytosis in various cell types^53, 55 54^, we propose that this is one mechanism through which residual viral proteins in the CNS^61^ contribute to persistence of neuroinflammation and neuronal damage despite suppressive ART.

Our cytokine multiplex analysis shows that, while iAst exposed to supernatants from HIV-infected iMg produce TNFα, this treatment is not sufficient to significantly induce the secretion of IL-6 by iAst (Supplementary Figure 3a). However, cocultures of HIV-infected iMg and iAst show a robust production of IL-6. iAst likely adopt a pro-inflammatory profile following exposure to HIV-infected iMg due to exposure to factors released from iMg such as cathepsin B and ATP following lysosomal exocytosis and begin secreting TNFα. In turn, iMg, which robustly express both TNFR-1 and TNFR-2 (Supplementary Figure 2b), respond to this signal by upregulating pro-inflammatory cytokine production, including IL-6 specifically, as TNFα can induce IL-6 production in various cell types^63, 64^. We speculate that this may be a mechanism through which these pro-inflammatory cytokines are produced at higher levels in PLWH, potentially contributing to damaging neuroinflammation and inflammatory pathology^8, 12^.

This pro-inflammatory signaling axis elucidated in our model has implications for various phenomena observed in PLWH. CSF viral escape, i.e., presence of detectable HIV RNA in the CSF despite suppressive ART with either undetectable or significantly reduced viral RNA in peripheral circulation, is observed in 5-15% of PLWH^19, 65^. Various explanations have been proposed for CSF viral escape, such as poor penetrance of ART drugs into the CNS, development of escape mutations that confer drug resistance to novel HIV strains, and suboptimal patient adherence to prescribed ART. Our results suggest another potential mechanism: dysregulated lysosomal trafficking in microglia harboring HIV leads to the activation of neighboring astrocytes into a reactive, cytokine-producing state. Subsequently, TNFα or other NF-kB activating cytokines produced by reactive astrocytes promote transcription from the integrated HIV genome, leading to viral replication. This may lead to a positive feedback loop as more microglia are infected and begin experiencing dysregulated lysosomal flux. Evidence for this potential mechanism also comes from recent studies reporting that CSF viral escape is associated with both increased neuroinflammation^66^ and astrocytic activation^67^. Future studies, such as single-cell RNA sequencing of brain tissue of SIV-infected rhesus macaques and tissue samples from PLWH experiencing CSF viral escape, are warranted to confirm and further elucidate the role of microglial lysosomal distress, astrogliosis, and NF-kB signaling in HIV-infected microglia as a potential signaling axis contributing to the persistence of neuroinflammation and neuronal injury despite suppressive ART.

A recent study has shown that damaged neurons induce HIV expression in previously latently infected microglia^68^, indicating a mechanism through which neuronal injury itself may lead to viral replication and inflammation, creating a positive feedback loop that in turn may worsen neuronal health and NCI^69^. Our data suggest a comparable mechanism through which reactive astrocytes rather than damaged neurons modulate HIV replication dynamics in microglia, driving increased replication via production of pro-inflammatory cytokines such as TNFα. Therefore, in line with many previous studies, our findings support the potential therapeutic role of anti-inflammatory agents to mitigate the incidence and severity of HIV-associated NCI in PLWH. Treatments designed to target reactive astrocytes may prove to be more feasible than those aimed at the rare HIV-infected microglia in ART-suppressed individuals, as astrocytes are a much larger population in the CNS.

Finally, our data showing dysregulated lysosomal positioning following HIV infection in microglia and associated lysosomal exocytosis, indicated by an increase in extracellular cathepsin B, suggest that lysosomal release may be involved in the propagation of a neuroinflammatory cascade in PLWH either as a legacy effect produced during acute infection or as a result of residual viral protein levels or low-level viral replication that may continue to occur despite ART. Accordingly, small molecules that can reduce lysosomal exocytosis and associated inflammation, such as the cannabinoid receptor 2 agonist JWH-133^53^, may be considered for adjunctive therapies to reduce HIV-associated NCI.

## Figure Legends

**Supplementary Figure 1:**
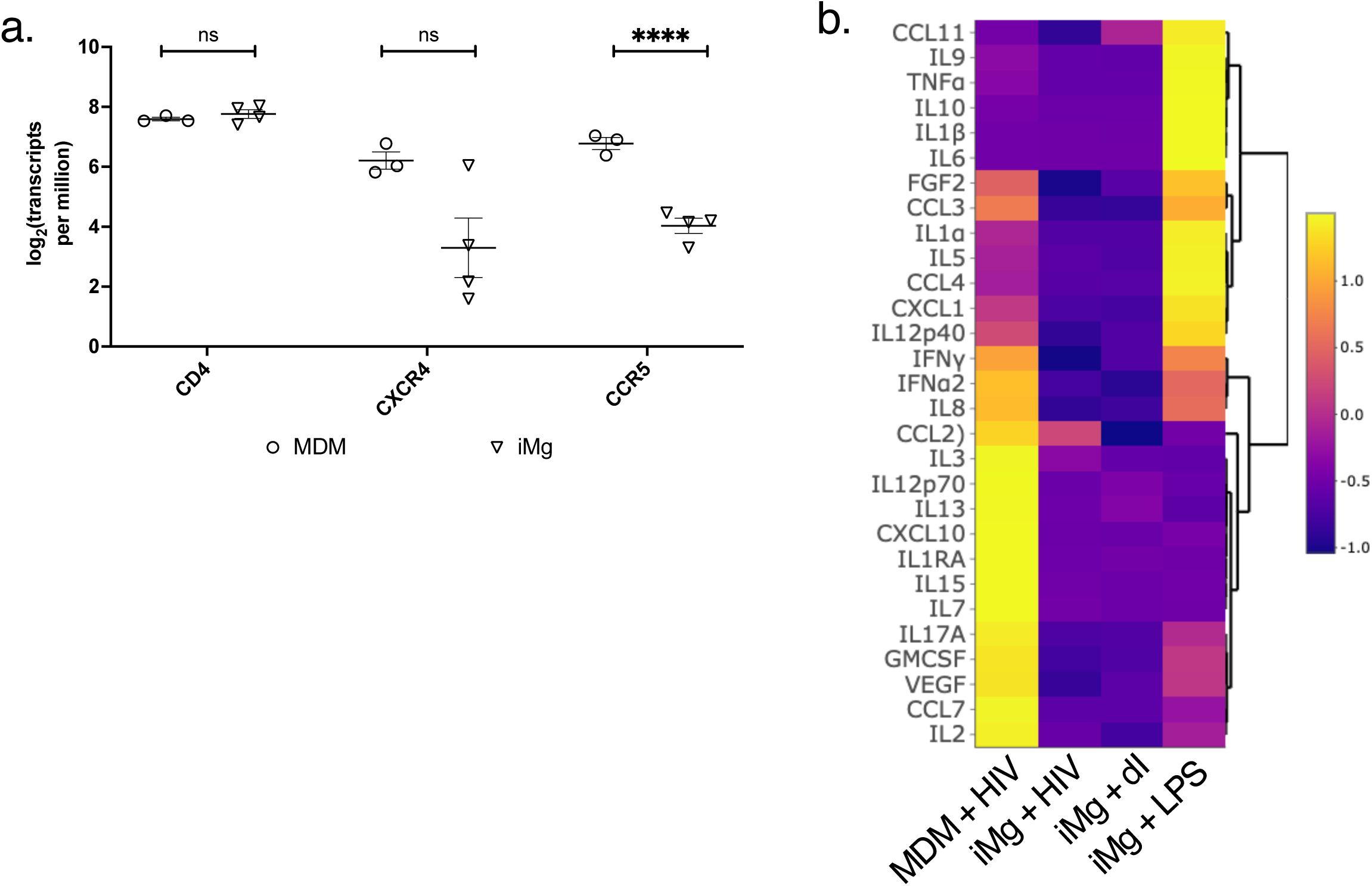
(a). Expression of indicated HIV-entry related genes in iMg at 11 DIV compared to expression in MDM in log transformed transcripts per million (b). Heatmap indicating relative levels of indicated cytokines and chemokines produced by HIV-infected MDM (left) or iMg (middle left) compared to mock-infected iMg (middle right) or iMg treated with 100 ng/mL LPS (right) at the protein level. Data are condensed from 2(MDM) to 5 (iMg) biological replicates. **** = p < 0.0001. Multiple paired T-tests.

**Supplementary Figure 2:**
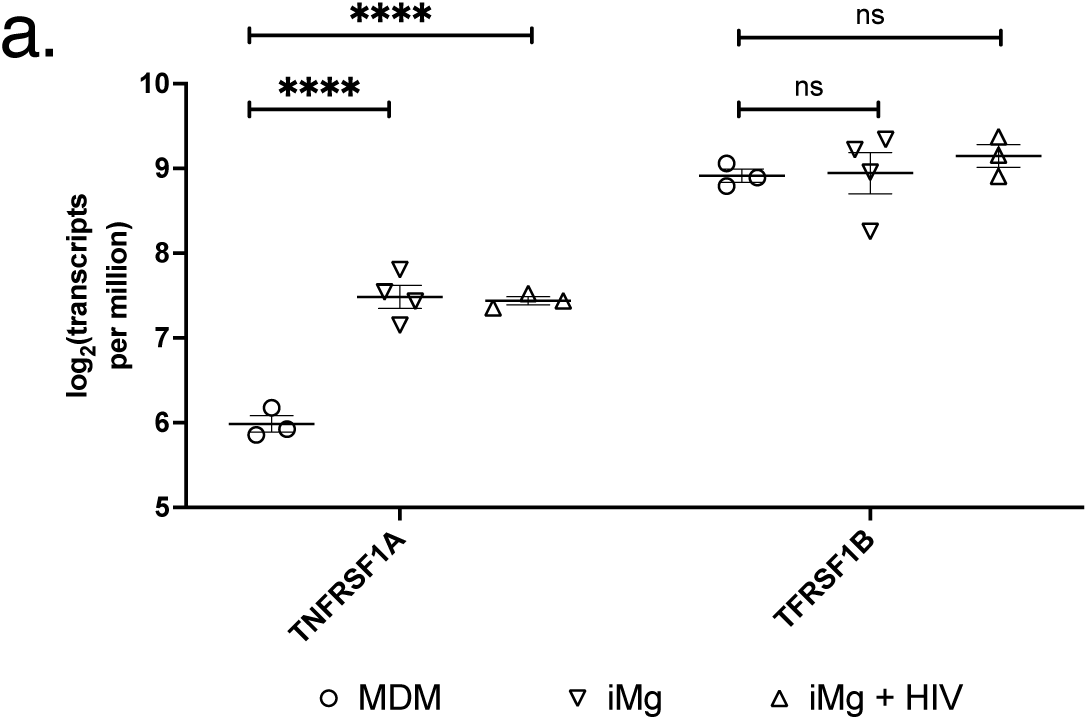
(a). Expression of indicated TNFɑ receptor genes in MDM and iMg with or without infection with HIV Jago at 50 ng/mL in log transformed transcripts per million. **** = p < 0.0001. Multiple paired T-tests.

**Supplementary Figure 3:**
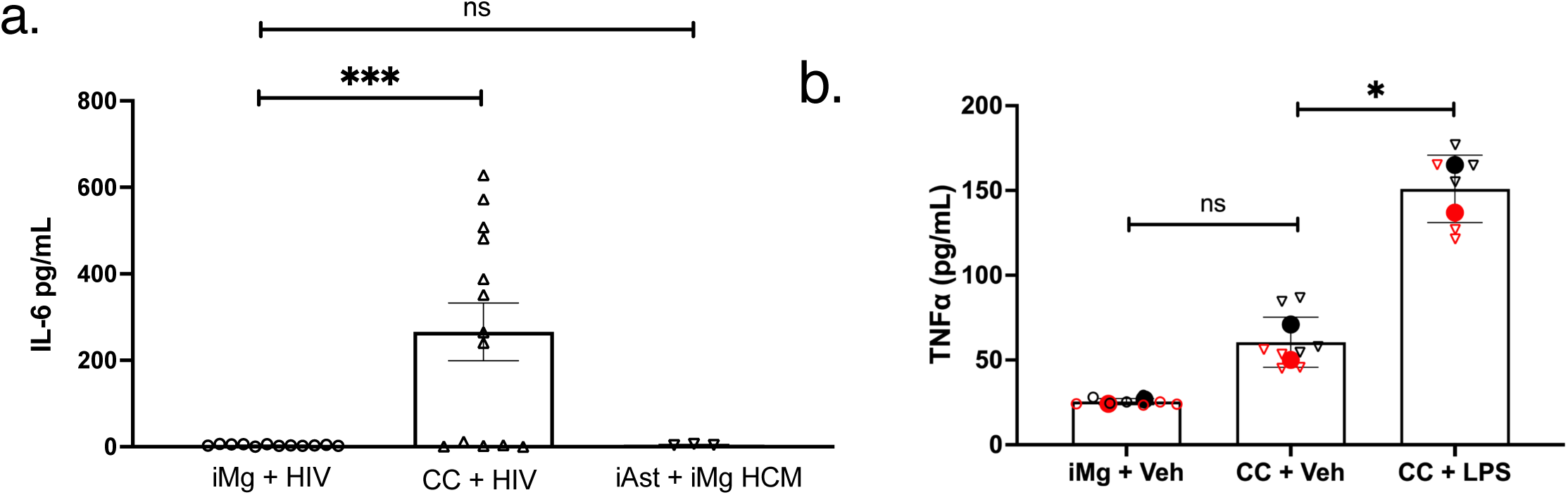
(a). Raw pg/mL values measured by biorad Luminex of IL-6 produced by HIV-infected iMg monocultures (circles), HIV-infected iMg + iAst coclutures (triangles), or iAst treated with conditioned media from HIV-infected iMg (inverted triangles). Data include 3 (iAst) or 5 (iMg, CC) biological replicates. (b). TNFɑ levels measured by alphalisa in iMg monocultures or iMg/iAst cocultures treated with vehicle (PBS) or 100 ng LPS. Technical replicates within a group of biological replicates are indicated by distinct colors, and averages from a given biological replicate are expressed as the overlaid solid-colored symbols. Data include 2 biological replicates. * = p < 0.05, *** = p< 0.001. Multiple paired T-tests(a), One-Way ANOVA(b).

**Supplementary Figure 4:**
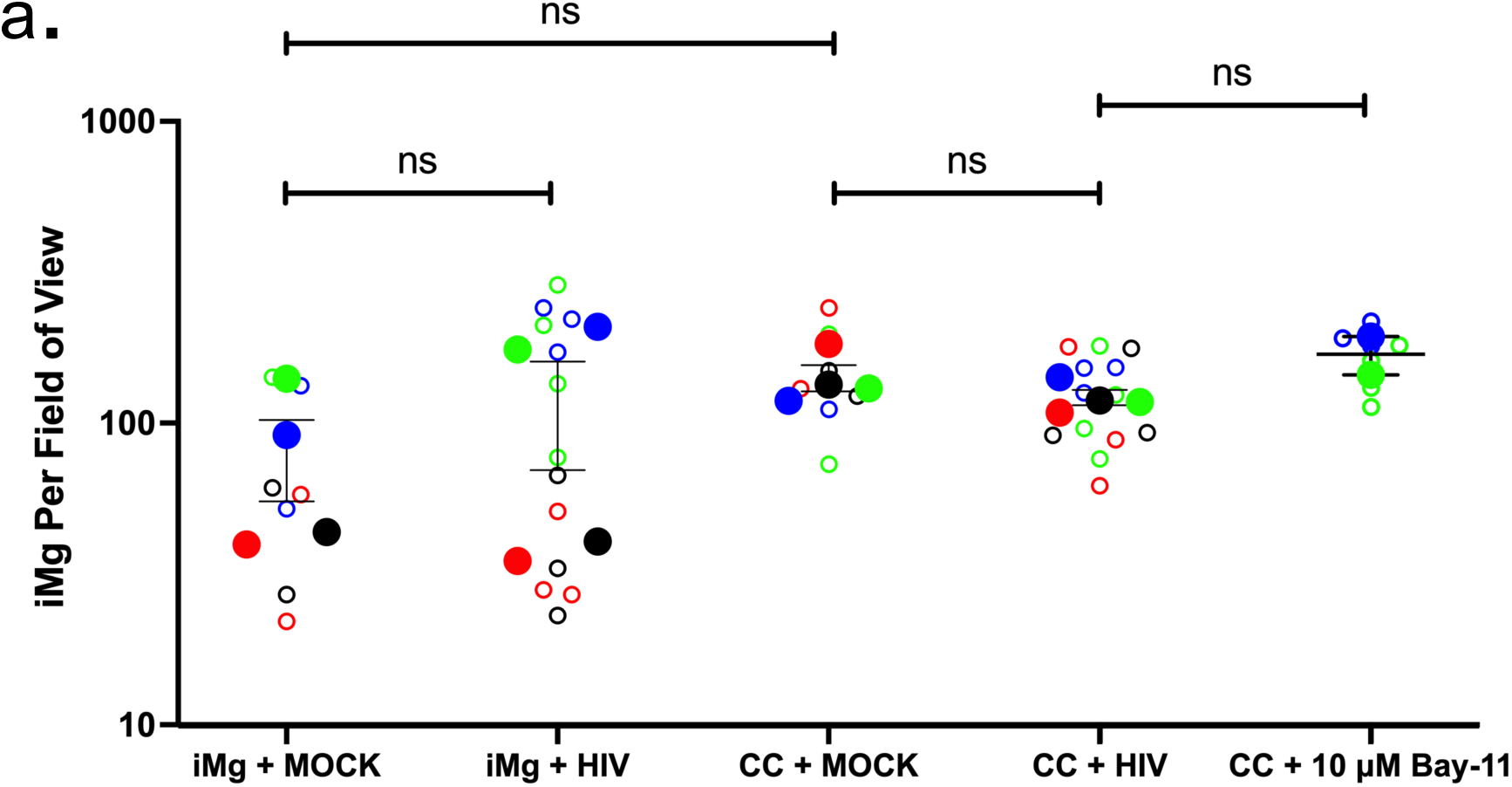
(a). Manual counts of nucleated IBA1-positive microglia per tile-scan via widefield microscopy at 40X in the indicated culture conditions * = p< 0.05. Multiple paired T-tests.

**Supplementary Figure 5:**
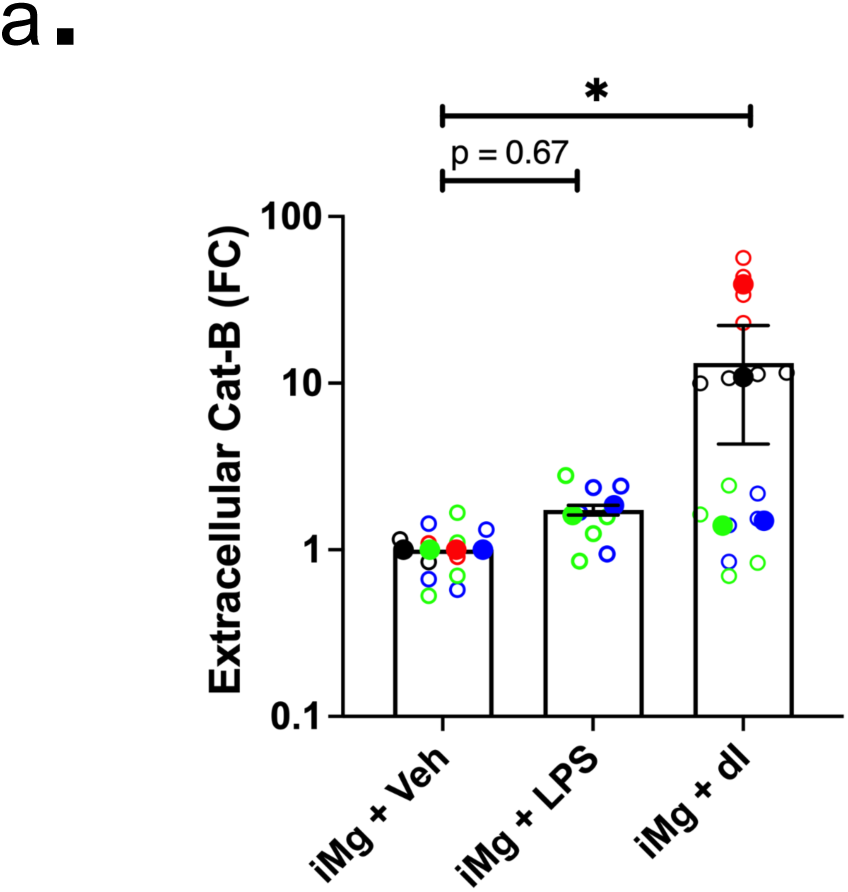
(a). Cathepsin B measured in culture supernatants from indicated groups at PID 12. Data are condensed from four biological replicates and plotted as fold-change vs the negative control group, iMg + Veh. * = p < 0.05, **** = p < 0.0001. Mann-Whitney test.

## Acknowledgments

We would like to thank all the members of the Gaskill lab and myeloid working group for invaluable discussion on the data. We would like to thank the donors who provided monocytes to the University of Pennsylvania Human Immunology Core. The following reagent was obtained through the NIH HIV Reagent Program, NIAID, NIH: Monoclonal Anti-Human Immunodeficiency Virus Type 1 (HIV-1) p24 Gag protein, Clone AG3.0 (produced *in vitro*), HRP-20068, contributed by Creative Biolabs.

## Author contributions

JG, CAE, and KLJS contributed study conceptualization and methodology. JG, JP, SS, EB, JS contributed experimental design, data acquisition, and data analysis. JG contributed original draft preparation. All authors contributed to writing, review, and editing. JG, PJG, CAE, and KJS contributed funding. All authors have read and agreed to the submitted version of the manuscript.

## Data availability statement

All sequencing data are available in the NCBI GEO database (https://www.ncbi.nlm.nih.gov/geo/) with the accession number GSE143685. Other data used and/or analyzed during the current study are available from the corresponding author on reasonable request.

## Funding

This research was funded by the National Institutes of Health grant no. T32 (JG), F31 MH131486(JG), R01DA057337(PJG), R61DA058501 (PJG), R21MH129193 (CAE), R01DA049514 and R01DA052826 (KLJS), and P30AI045008 (KJS, CAE).

## Notes

https://www.ncbi.nlm.nih.gov/geo/query/acc.cgi?acc=GSE143685

### Competing Interest Statement

The authors have declared no competing interest.

### Summary of Updates

Correcting typos in introduction and methods and improving a figure legend in Fig S1

